# The genomic landscape of metallic color variation in ground beetles

**DOI:** 10.1101/2023.09.25.559374

**Authors:** Yi-Ming Weng, David H. Kavanaugh, Bryan Rubio-Perez, Jad Salman, Mikhail A. Kats, Sean D. Schoville

**Author notes:** Corresponding author: Yi-Ming Weng, **Email:**.

## Abstract

The metallic color variation of beetles is a spectacular feature that has inspired diverse human cultures. However, little is known about the genetic basis of this trait or its ecological importance. In this study, we characterize the geographical distribution, optical mechanism, genetic basis, and ecological and evolutionary importance of metallic color variation in the *Nebria ingens* complex, an alpine ground beetle in the Sierra Nevada, California. We find that elytral color varies continuously across two allopatric species (from black *N. ingens* to green *N. riversi*), with hybrid populations showing intermediate coloration, and we demonstrate that the metallic color is generated from multilayer reflectors in the epicuticle of the elytra. By applying association mapping in natural populations (wild-GWAS) using high-density genotype variants, we identify five promising candidate genes covarying with metallic variation, with known roles in cuticle formation and pigmentation pathways. This finding, together with a significant correlation between color variation and water availability, suggests that metallic variation evolves as a local adaptation to environmental variation in the *N. ingens* complex.

## Introduction

The impression created by bright metallic coloration in beetle wings is so stunning that diverse human cultures have used them in art and jewelry (‘beetlewing art’) for thousands of years (Ratcliffe, 2006; Rivers et al., 2003). This remarkable phenotypic trait is all the more impressive when its tremendous variation among beetles (order Coleoptera), a biodiverse group including more than 500,000 species, is recognized to have repeatedly evolved (Seago et al., 2009). The optical mechanism of metallic color in insects is well known, generated by the internal structure of the exoskeleton (Andersen, 2009). In Coleoptera, the most prevalent internal structure is a multilayered reflector in the epicuticle or exocuticle, where the layers with alternating refractive index reflect light in a specific range of wavelengths (Barry et al., 2020; Seago et al., 2009). For species exhibiting multilayered reflectors, variations may involve differences in saturation (chroma), hue, or both. The thickness of the alternating layers primarily governs the hue, with thicker layers producing longer peak wavelengths and thinner layers resulting in shorter ones (Kurachi et al., 2002). Meanwhile, saturation can be influenced by the number of layers or the overall tanning level of the cuticle. As a result of the discovery of the optical mechanism of metallic colors in insects, these microstructures have become an important resource for bioinspired materials (Biró et al., 2010; Biró & Vigneron, 2011; Colusso et al., 2017).

Although metallic coloration is known to be polymorphic in numerous beetle species (Kozlov et al., 2022; Strickland et al., 2019), little is known about the genetic basis of this phenotypic trait. Compared to beetles, the genetic basis of wing coloration in butterflies has been more extensively studied and could provide hypotheses for testing the roles of similar genes in regulating multilayer reflectors in beetles. For example, transcription factors such as *optix, Antennapedia, aristaless1, bric-a-brac* (*bab*), and *spalt* are involved in various pigmentation pathways and in regulating the photonic microstructures on butterfly scales (Ficarrotta et al., 2022; Livraghi et al., 2025; Mazo-Vargas et al., 2022; Prakash et al., 2022; Reed et al., 2020; Thayer et al., 2020; Zhang et al., 2017). These transcription factors may also regulate the synthesis, transport, or deposition of melanized layers in the beetle cuticle. On the other hand, genes involved in the tyrosine-mediated tanning pathway such as *ebony, tan, arylalkylamine N-acetyltransferase* (*aaNAT*), *laccase2, yellow, pale* (*tyrosine hydroxylase*), *Dopa decarboxylase* (*DDC*), along with potential transcription factors such as *pannier* (*pnr*) and *POU domain motif 3* (*pdm3*), are responsible for the production and distribution of melanin pigments and could therefore contribute to patterning the multilayer reflector (Arakane et al., 2016a; Niimi & Ando, 2021).

Since studies of diverse beetle species have shown that metallic color variation is typically additively determined by a small number of genetic loci (Favila et al., 2000; Fujiyama & Arimoto, 1988; Gotoh & Lavine, 2014; Mossakowski & Paarmann, 2014a; Vasconcellos-Neto, 1988), and only infrequently dependent on genotype-environment interactions (Davis et al., 2008), it should be possible to identify the genes underlying the metallic color variation by association approaches (Wellenreuther & Hansson, 2016).

In this study, we aim to investigate the genetic basis of metallic color from the multilayer reflector in the cuticle of alpine ground beetles, the *Nebria ingens* species complex, by using whole genome sequence data and association mapping in natural populations (wild-GWAS). The *N. ingens* species complex is endemic to the Sierra Nevada in California, where it is found in the high elevation nival zone that is covered by snow throughout most of the year. Two species with divergent color in their elytra (appearing metallic green or black to human perception), *N. ingens* Horn and *N. riversi* Van Dyke, are geographically isolated, but form a broad admixture zone of intermediately colored individuals (Schoville et al., 2012; Weng et al., 2021) (**Figure 1**). The morphologically variable and genetically admixed populations make this species complex a great study system to apply wild-GWAS approaches (Buerkle & Lexer, 2008; Lindtke et al., 2013). Replicate admixed populations disrupt linkage disequilibrium across the genome, increasing power to isolate causative loci underlying the divergent phenotypes of the two species. Furthermore, the nearly continuous color variation improves the statistical power of association analyses. Leveraging this system, we identify candidate for the genetic basis of metallic color in beetles which could be tested in the future expression regulation or gene editing approaches.

**Figure 1.**
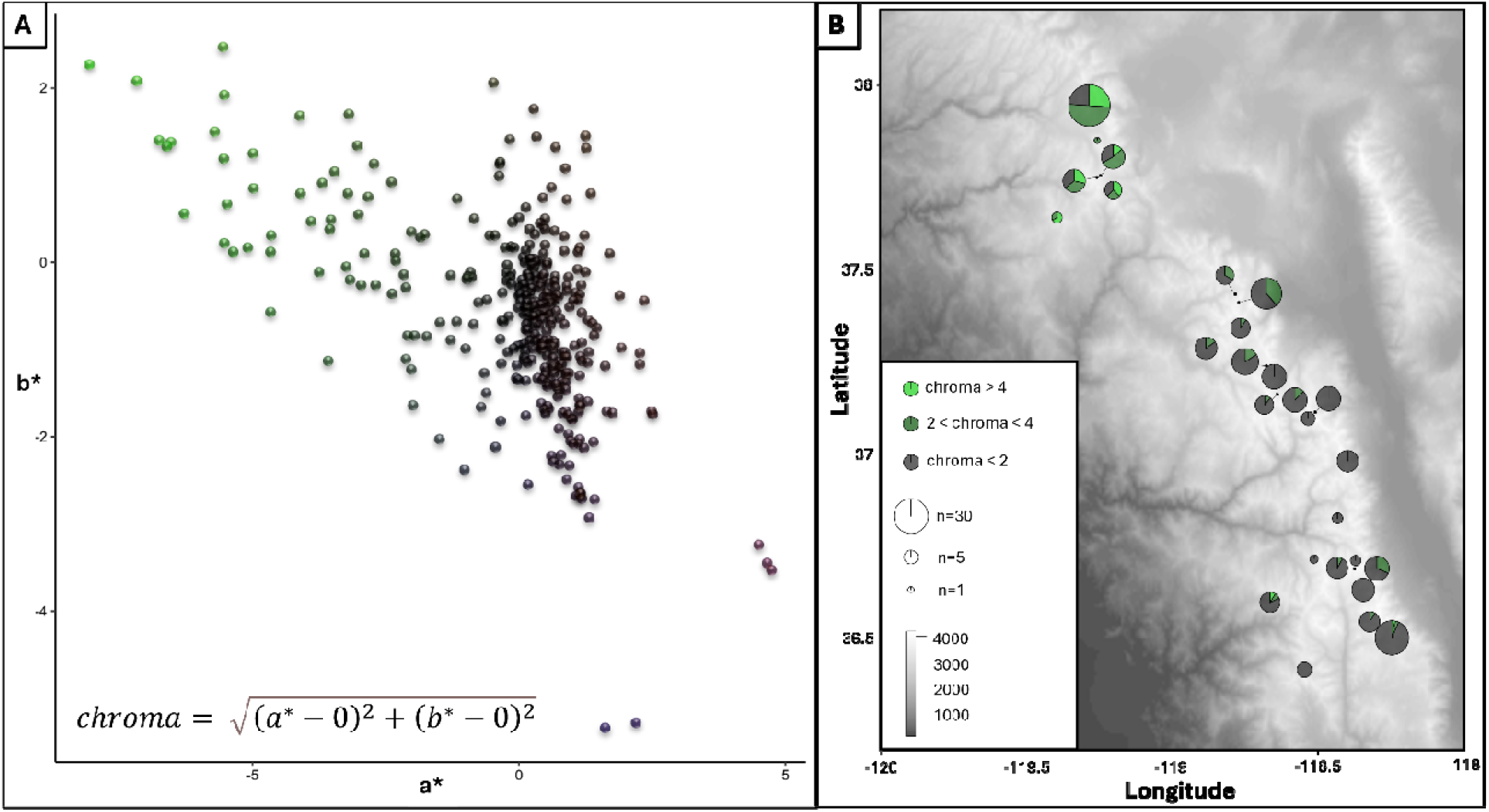
Color variation of the elytra in the *Nebria ingens* species complex. (A) Quantification of color using the a^D^ (green-red axis) and b^D^ (blue-yellow axis) color space, where chroma is defined as the distance of each point from the achromatic center (0,0). (B) Map showing the geographic distribution of color variation across sampling sites: the northernmost six populations correspond to *N. riversi*, the southernmost eleven populations to *N. ingens*, and the intermediate populations represent admixed groups.

## Materials and Methods

### Animal, genotyping, and color variation

Adult beetles of the *N. ingens* complex (n=383) were collected from 27 populations, encompassing the entire distribution range of the species complex (**Figure 1**). Whole-genome resequencing data from 374 beetles were generated in our previous study using the GATK variant calling pipeline (Van der Auwera & O’Connor, 2020; Weng et al., 2024). After removing samples lacking color phenotypic data and loci with minor allele frequency less than 0.05 using vcftools v0.1.16 (Danecek et al., 2011), we performed GWAS using a genotype matrix with 355 beetles and 526,717 SNP+indel variants. To quantify optical variation, we measured the reflectance spectra from the elytral surface of 368 beetles using a hyperspectral camera (Specim IQ). The average spectrum was measured from each pixel in the brightest elytral region (8-600 pixels, median=190 pixels) to represent the individual reflectance spectrum. The mean reflectance spectrum was then used to calculate the CIELAB color for each elytron (Robertson, 1977) using the CIE standard illuminant D65 with a custom MATLAB script. Chroma was calculated as the distance from the neutral gray point (0,0) in the a*b* plane to represent the color saturation of the elytra.

### Cuticular structure of the metallic hue

The elytra of three beetles, representing *N. ingens, N. riversi*, and a genetically admixed hybrid lineage, were used to examine the cuticular microstructure with transmission electron microscopy (TEM). We used a FEI Tecnai T12 microscope in the School of Medicine and Public Health at UW-Madison. The samples were stored in -20°C with 95% ethanol before use. The images with clear structure were selected to measure the thickness of the epicuticle. We took 13-30 vertical transects to repeat measurements of thickness for each specimen using ImageJ (Abràmoff et al., 2004). Counts and thickness of layers were used to predict the reflectance spectra of each specimen using a multilayer reflectance calculator provided by Filmetrics^®^ Corporation, with the refractive indices of electron dense and electron lucent being 1.7 and 1.4, respectively (**Figure 2**). These indices are consistent across different beetle taxa (Bernard & Miller, 1968; Kurachi et al., 2002). The calculated reflectance spectrum was subsequently used to compare with the observed spectra from the same individual beetle, except for the green elytrum for which we compared the calculated spectrum to other bright green beetles from the same population, to verify the source of the metallic color (**Figure 2**).

**Figure 2.**
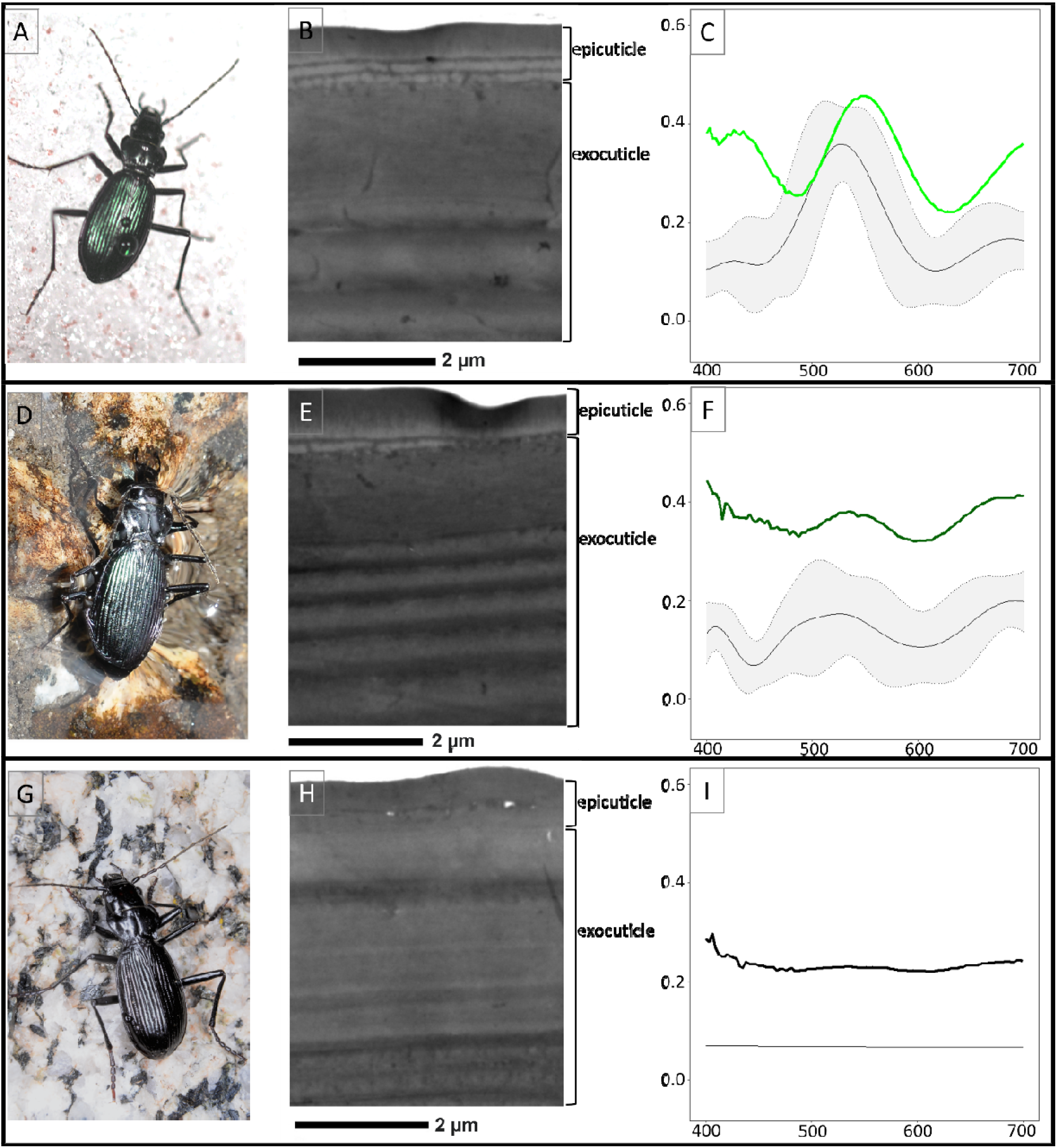
Comparisons of modeled spectra calculated from measurements of the multilayer reflectors in the epicuticle of individuals with observed reflectance spectra in different elytral colors (A-C: green, D-F: dark green, G-I: black). The TEM images demonstrate the pattern of multilayer reflectors in the epicuticle. In each spectrum plot, the thin lines represent the means of modeled spectra from different observation points (see Supplemental methods) and the grey area shows the 95% confidence interval from measurements of different vertical transects, and the measured spectra are denoted with colored thick lines.

### Genome-wide association mapping

To identify SNP and indel markers associated with the color phenotype, we performed GWAS using latent factor mixed models (LFMM) and partial redundancy analysis (pRDA) implemented using R package lfmm and vegan, respectively (Caye et al., 2019; Oksanen et al., 2025). In LFMM analysis, four latent factors were specified to account for population structure, which includes a pattern of isolation by distance across seven ancestral population clusters (Weng, Kavanaugh, et al., 2021). The regularized least squares estimate of causal variants were obtained using both ridge and lasso regression. To reduce the false-discovery rate (FDR), we first calibrated the *p* values with genomic inflation factors (GIF), then adjusted the calibrated *p* values using the Benjamini-Hochberg multiple-testing procedure. Variants with the adjusted *p* value less than 0.01 in ridge or lasso regression were considered candidates of LFMM. In pRDA analysis, genotype data was first transformed into plink format (.raw) using vcftools v0.1.16 (Danecek et al., 2011). To adjust the linear effects of chroma on genotype while accounting for the variation from the population structure, the first four PC axes calculated with SNPRelate were included as conditioning variables (Zheng & Zheng, 2013). Variants falling outside of four standard deviations of first RDA axis loading are considered candidate variants for chroma. The intersection of candidate variants from the two analyses were used to find the intersecting genes from the published genome using bedtools v2.28 (Quinlan, 2014; Weng, Francoeur, et al., 2021). Gene intersection was defined by variants falling into the genetic region and their 2 kilobase upstream (5’) flanking region, as regulatory elements could be located thousands of base pairs from the 5’ flanking region. Finally, we assessed to annotated function of the candidate genes by referencing their best blast hit against NCBI nr database and InterPro function using InterProScan (Blum et al., 2025; Camacho et al., 2009; Jones et al., 2014), and compared candidate genes with our previously published genes with signal of selective sweeps from OmegaPlus to identify overlapping genes potentially involved in both color variation and evolutionary adaptation (Alachiotis et al., 2012; Weng et al., 2024).

### Testing for environmental selection on color variation

Because color variation is a key diagnostic character separating two species in the *N. ingens* complex, it may be associated with adaptation to local environments. We investigated the correlation between color variation, water loss rate of beetles, and the snow water equivalent (SWE) in the habitats of the sampled populations. To quantify the water loss rate, 87 beetles were placed individually in a tube in a sealed container, which contained lithium chloride (LiCl) as a desiccant to maintain a relative humidity between 20 and 30 percent. The temperature was controlled at 5°C during the experiment, which is similar to the natural foraging temperature in the field. The water loss rate was calculated by weighing beetles before and after 24 hours of desiccation. Habitat SWE data were extracted from published Sierra Nevada snow analysis results using the R package raster v3.6-3, with further methodological details provided in our previous study (Margulis et al., 2016; Weng et al., 2024). Spearman’s rank correlation coefficient (ρ) was used to assess the correlation since the data were non-normal in distribution.

## Results

### Continuous variation and the mechanism controlling metallic color

We examined 368 beetles from 27 populations across the distribution of the species complex. In general, the reflectance spectra of green beetle elytra have a peak of 500 to 600 nm wavelength, while beetles with black elytra have a lower reflectance amplitude and loss of the characteristic reflectance peak resulting in the green color (**Figure 2**). By converting the reflectance spectra to the CIELAB color space (L*a*b*) using the CIE standard illuminant D65, we found that many, but not all beetles identified as *N. riversi* reflect a green hue, while most beetles identified as *N. ingens* or as hybrids reflect a dark green to black hue (**Figure 1**). The color variation is continuously distributed and thus likely to be a quantitative trait encoded by multiple genes.

From the elytral cross section, TEM images of green, dark green, and black elytra demonstrate variation in the inner epicuticle (cuticulin layer), which is most pronounced in the green elytra (**Figure 2**). There are 1-3 electron dense layers 50-100 nm thick (darker layers in the images due to reduced permeability for the electron beam), alternating with similar-width electron lucent layers, serving as a multilayered reflector (**Figure S1**). In the dark green elytra, only one electron dense layer is found, and no clear multilayer reflector was observed in the black elytra throughout the examined regions. Calculated spectra based on the thickness of the multilayer reflectors generally matches the visual color and observed spectra, with more alternating layers producing a stronger amplitude of light at a peak wavelength between 500 and 600 nm, roughly corresponding to the color green (**Figure 2**).

### Color associated genes and their roles in color determination

Among the 526,717 scanned variants, 892 (0.17%) passed the adjusted *p*-value threshold at 0.01 in ridge or lasso regression models and 573 (0.11%) fall outside of 4^th^ standard deviation of first RDA axis loading distribution, where 195 (0.037%) variants are found shared in both candidate lists and considered candidate variants. These candidate variants intersect 127 genes including 65 genes with functional annotations (**Table S1**) and five with known functions in different pathways of insect cuticle formation and pigmentation (**Figure 3**). These include *yellow-e* and *pdm3* in the tyrosine-mediated tanning pathway, *aristaless* in the ommochrome pathway, *chico* (encoding insulin receptor substrate (IRS) protein) in the Insulin/IGF signaling pathways through PI3K/Akt, and a gene encoding Pyrokinin-1 receptors-like protein (*PK1*-like), a type of G-protein coupled receptor in the pyrokinin signaling pathway (Matsumoto et al., 1990; Shakhmantsir et al., 2014). All five genes have intersecting variants within intragenic regions, and the variants in *pdm3* and *chico* are in the coding region. Finally, the candidate genes *chico, PK1*-like, and small G protein signaling modulator 3 (*GPSM3*) homolog were previously identified in genome-wide scans for selective sweeps.

**Figure 3.**
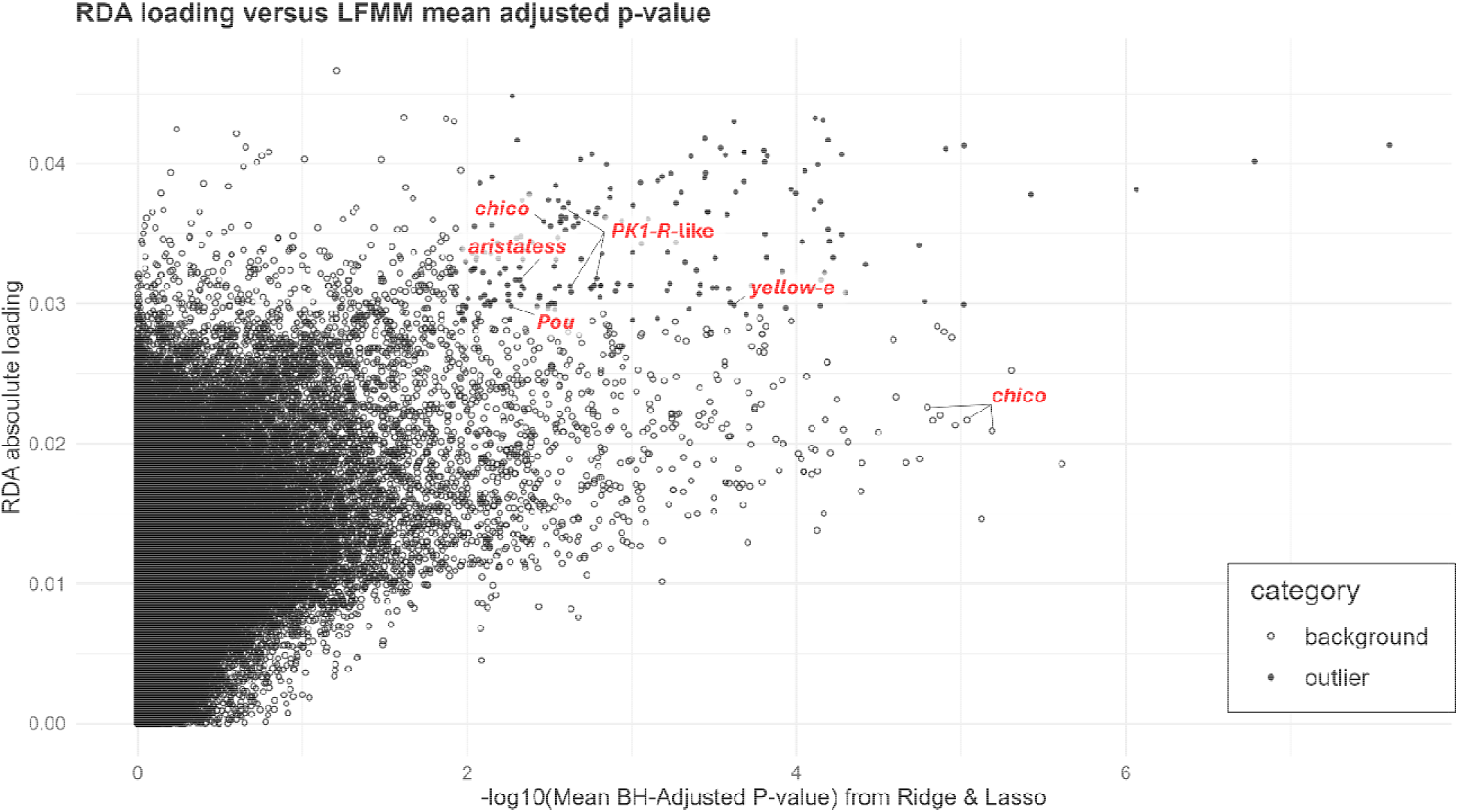
The scatter plot with mean BH-adjusted p-value of LFMM (x axis) and RDA absolute loading (y axis). Each dot represents a variant where the close dots are the outlier (candidate) variants defined by LFMM BH-adjusted p-value of ridge or lasso less than 0.01, and RDA loading falling outside of 4^th^ standard deviation.

### Correlation between color variation, water loss rate, and SWE

To test the physiological and ecological function of metallic color, we performed pairwise correlation analysis using chroma, water loss rate, and seasonal water availability in the nival zone (represented by measurements of snow water equivalent in September, SWE) (**Figure 4**). The results showed a significant positive correlation between beetle color and SWE (ρ=0.45, 95% CI=0.35 to 0.53; *p*= 2.2e-16), with higher color chroma occurring in more snow-rich habitats, and a significant positive correlation between water loss rate and SWE (ρ=0.26, 95% CI=0.06 to 0.44; *p*=0.014). We also found a positive trend between color variation and water loss rate but that was not significant (ρ=0.21, 95% CI=-0.06 to 0.45; *p*=0.12).

**Figure 4.**
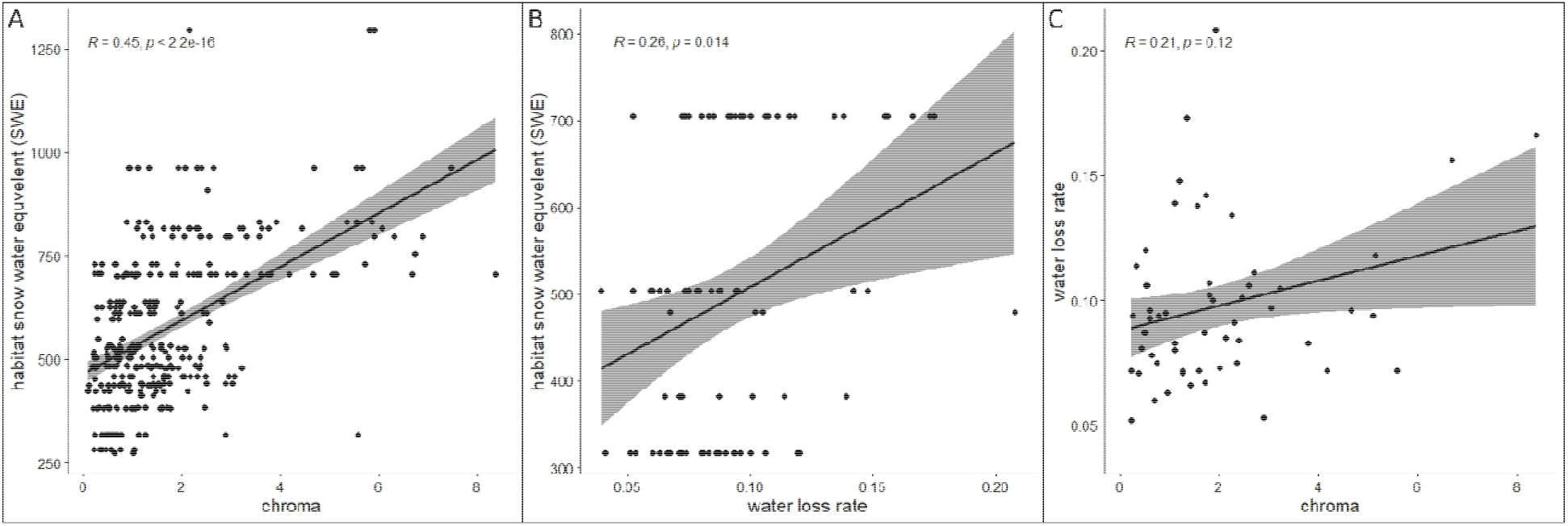
The correlations among elytral color (chroma), water loss rate, and mean September snow water equivalent (SWE) in habitat of the *N. ingens* complex. From left to right: (A) SWE vs. chroma; (B) SWE vs. water loss rate, and (C) water loss rate vs. chroma. The correlations were calculated using Spearman’s rank correlation coefficient, and the grey areas indicate the 95% confidence interval of the estimate.

## Discussion

Despite a long-standing interest in the metallic color of insects, distinct knowledge gaps in our understanding of color variation remain. Beyond the lack of knowledge for the most biodiverse animal group, the beetles, gaps include the nebulous genetic basis of metallic color variation. By applying wild-GWAS to an alpine ground beetle species complex with admixed populations, we found promising candidate genes that are statistically associated with the metallic variation generated from multilayer reflectors in the epicuticle and functionally involved in cuticular structure and melanization. These include genes from several known pathways involved in butterfly color patterns.

### Genetic basis of the metallic color in the N. ingens complex

Color variation can be categorized by optical mechanisms which vary across insect lineages. For instance, in Coleoptera, iridescent color generated by three-dimensional photonic structures has been found in weevils and longhorn beetles, whereas multilayer reflectors are widespread in distantly related lineages of jewel beetles (Buprestidae), ground beetles (Carabidae), leaf beetles (Chrysomelidae), and scarab beetles (Scarabaeidae) (Seago et al., 2009, 2019). Although structural layering may be common to the ancestors of beetles, structural coloration has been repeatedly gained, lost and modified (Seago et al., 2019). Yet very little is known about how genetically complex this phenotypic variation is across beetles.

In this study, we found that a multilayer reflector is responsible for the metallic color variation of the *N. ingens* complex. This is directly supported by the presence of the layers with alternating refractive index in the epicuticle, with their estimated reflectance spectra matching the elytral colors and the observed spectra (**Figure 2**). Such resemblance of the calculated and observed reflectance spectra has been used to determine the optical mechanism of metallic color in many other insects (Bermúdez-Ureña et al., 2020; Kurachi et al., 2002; J. Lee et al., 2020; Luna et al., 2013; Noyes et al., 2007; Parker et al., 1998; Pasteels et al., 2016). In the *N. ingens* complex, the amplitude of the peak between 500 and 600 nm in the reflectance spectrum is directly correlated with the number of alternating layers in the epicuticle, representing a form of saturation variation characterized by changes in amplitude at specific wavelengths caused by differing numbers of layers with consistent thickness. It is structurally distinct from phenotypes where the reflectance peak shifts across wavelengths due to differences in layer thickness and therefore should be regarded as a separate category of color variation. In nature, this type of saturation variation seen in the *N. ingens* complex appears to be less common but has been reported in different beetle families (Gotoh & Lavine, 2014; Mossakowski & Paarmann, 2014b; Stanbrook et al., 2021).

Knowing that the metallic color variation in the *N. ingens* complex is regulated by multilayer reflectors, we conducted a wild-GWAS to identify genetic variants associated with this color variation. We discovered 65 genes containing variants that significantly correlate with the chroma of the elytra, including five genes with known roles in cuticle pigmentation processes (**Figure 3**). Among these five genes, *yellow-e* and *pdm3* in the tyrosine-mediated tanning pathway are most likely to contribute to the color variation in the *N. ingens* complex, as the multilayer reflectors are primarily composed of alternating layers with different degree of melanin deposition (Hamza & Zahran, 2023; Onelli et al., 2017; Parker et al., 1998). Yellow protein (dopachrome conversion enzyme) from the *yellow* gene family advances melanin biosynthesis, a vital biochemical pathway altering the insect cuticle profile by converting tyrosine into pigments (e.g., melanin and dopamine-melanin) (Arakane et al., 2016b; Noh et al., 2016). This gene has been identified as a key mediator of the melanin biosynthesis pathway across multiple insect species (Gompel et al., 2005; Li & Christensen, 2011; Noh et al., 2015; Zhang, Martin, et al., 2017). On the other hand, the transcription factor *pdm3* is known for its role in regulating neural development, but it has also been identified as a repressor of abdominal pigmentation in *Drosophila melanogaster*, potentially through co-expression with other melanin biosynthesis transcription factors such as *Abdominal-B* and *doublesex* (David et al., 2025; Hughes et al., 2023; Kim & Cho, 2020; Rogers et al., 2014; Yassin et al., 2016). However, the function of *pdm3* orthologs in beetle (Coleoptera) species remains unexplored, highlighting the need for further studies that investigate its role in beetle cuticle pigmentation. In contrast, *chico* and *PK1-like* genes are indirectly involved in cuticle pigmentation through the Insulin/IGF and pyrokinin signaling pathways, respectively (Matsumoto et al., 1990; Shakhmantsir et al., 2014). Although their involvement is indirect, signals of positive selection from the selective sweeps have been previously identified in these genes in the *N. ingens* complex, underscoring the importance of further investigating their roles in the pigmentation of beetle elytra (Weng et al., 2024). Finally, *aristaless*, a transcription factor known to influence appendage formation and scale color in butterflies, has been identified as a candidate gene (Bayala et al., 2023). However, *aristaless* is involved in ommochrome pigment production through the ommochrome pathway.

Since melanin is the dominant pigment in insect cuticles, we consider *aristaless* the least likely candidate among the five. In summary, we identified five genes linked to beetle elytra color, with *yellow-e* and *pdm3* most likely influencing melanin-based pigmentation. *Chico* and *PK1*-like show indirect roles and signs of selection, while *aristaless*, involved in a different pigment pathway, is the least likely candidate.

### The evolutionary and ecological role of metallic color in the <u>N. ingens</u> complex

The ecological function of metallic color has been considered an important characteristic in biotic interactions, both within and between species (Clusella-Trullas & Nielsen, 2020). In beetles, sexual selection, mimicry, camouflage, or aposematism are widely discussed functions for metallic color (Kjernsmo et al., 2022; Lee et al., 2018; Vulinec, 1997; Xu et al., 2010). But just how relevant biotic interactions are for the diversity of color patterns in beetles remains unclear (Ospina-Rozo et al., 2024; Schultz & Hadley, 1987). Other studies have shown that metallic color may instead result from adaptation to abiotic environmental factors (Davis et al., 2008; Keinath et al., 2020). In particular, pigmentation from the melanization pathway in the epidermal and serosal cuticle is thought to be associated with desiccation resistance (Bai et al., 2022; Farnesi et al., 2017; Wang et al., 2021). In the subgenus *Catonebria*, which includes the *N. ingens* complex, metallic color variation has been gained and lost many times with a general trend that metallic species occur in cooler and wetter regions compared to related species with darker, less or non-metallic cuticle. For example, among species of the *Nebria gebleri* species complex (Kavanaugh et al., 2021), those from more northern and coastal regions (northern Cascade Range and northern Rocky Mountains; namely, *N. gebleri* Dejean and *N. cascadensis* Kavanaugh) are more vividly metallic than those from the Sierra Nevada and other southern ranges in California (i.e., *N. albimontis* Kavanaugh, *N. rathvoni* LeConte, and *N. siskiyouensis* Kavanaugh). The same general pattern is seen among members of the *meanyi* species complex as well as in other closely related *Nebria* subgenera. To investigate this phenomenon, we explored the possibility that metallic variation might be associated with water availability in *N. ingens* habitats, with the expectation that the black individuals could have higher fitness in drier and warmer habitats. Evidence from other beetle species has shown a relationship between the *yellow* gene, cuticle melanization, and water retention ability (Noh et al., 2015). This leads to the hypothesis that the loss of metallic color, as a byproduct of melanin accumulation, is the result of adaptation to arid environments. Our results show correlations between chroma and habitat SWE, as well as between water loss rate and habitat SWE, indicating that less melanized beetles are more commonly found in habitats with lower snow covers (**Figure 4A & 4B**). Moreover, individuals inhabiting these drier environments tend to exhibit better water retention ability. However, the correlation between chroma and water loss rate is moderate and not statistically significant (**Figure 4C**), suggesting that additional factors could be contributing to variation in water retention ability.

### Study limitations and future directions

While our wild-GWAS approach successfully identified genetic variants associated with beetle elytra color variation, several limitations may have influenced the strength of the association signals. Despite efforts to minimize noise, challenges in accurately quantifying color, such as measurement errors due to varying specimen conditions, could have introduced variability into the data. Additionally, nearly half of the candidate genes predicted from the reference genome lack functional annotation, which means that important genes may have been overlooked in our analysis. Nonetheless, we identified five promising candidate genes that warrant further functional investigation to better understand their roles in producing the metallic coloration in beetles.

## Acknowledgments

We thank Filmetrics^®^ for granting permission to use the reflectance calculator. We thank the National Parks of the United States, including Yosemite, Sequoia and Kings Canyon National Parks, for providing permission to conduct research in the parks. This research was financially supported by the Valentine Eastern Sierra Reserve (UC Santa Barbara Natural Reserve System) through VESR Graduate Student Funding and the Ministry of Education of the Republic of China (Taiwan) through the Government Scholarship to Study Abroad to Y.M.W. We thank the PhD committee of Y.M.W., consisting of Daniel Young, Carol Lee, Prashant Sharma, and Jun Zhu, for providing suggestions regarding this study. The authors have no conflicts of interests to declare.

## Author Contributions

YMW, DHK, and SDS developed the original idea; YMW and SDS designed the experiments. The quantification of color variation was conducted by YMW, BRP, and JS and other analyses were performed by YMW. YMW and SDS wrote the first draft of the manuscript and DHK, BRP, JS, and MAK provided comments to improve the manuscript.

## Notes

### Competing Interest Statement

The authors have declared no competing interest.

### Summary of Updates

This revised manuscript includes revised analyses and results. Some analyses and resulted are removed from the previous version.

