## supplemental_Information for "The genomic landscape of metallic color variation in ground beetles"

**Supplemental methods**

*Genotypic data preparation for LFMM*

We used the single nucleotide polymorphism (SNP) data matrix generated from our previous study (Weng et al., 2024) as the genetic marker to perform GWAS using latent factor mixed models (LFMM). The SNP matrix containing 526,717 SNP+indel variants was originally called from 374 beetle genomes, with series of filtering preprocessed in the previous study and stored in the variant call format (vcf) file, including the removal of loci with minor allele frequency less than 0.05. In this study, we first subset the variant matrix to 355 beetle individuals with color variation data using SelectVariants function in GATK (Van der Auwera & O’Connor, 2020). The subset SNP matrix was subsequently converted to ped format using vcftools v0.1.16 for LFMM (Danecek et al., 2011).

*Quantification of Color Variation*

Metallic color variation of adult beetles varies as a reflection spectrum on the elytra. We measured reflectance using a hyperspectral camera (Specim IQ) to record every pixel in a dorsal image of each ethanol preserved specimen (n=368), comprised of a 512 × 512 pixel array. The spectral resolution of the image is 7 nm and the spectrum ranges from 400 to 1,000 nm, with 204 spectral wavelength points in this range. The images were then used to extract the refection spectrum by manually selecting the brightest region of the elytra (because the elytral surface is convex and depends on the location of the input light). Selected areas consisted of 200 pixels on average (8-600 pixels, median=190 pixels), and the mean reflectance was calculated from the selected pixels for each of the 204 spectral wavelength points. Finally, we used the mean reflection spectrum to calculate the CIE LAB color space for each elytron using CIE standard illuminant D65 (Robertson, 1977), in which we used axis a* (the axis of green-red) to represent the color of the elytral sample. We chose this color axis because it encompasses the most variation in elytral color.

*Cross-section TEM of elytra and calculated reflectance spectra*

From the cross-section TEM images of the green, dark green, and black elytra, we found multilayer reflectors embedded in the inner epicuticle (cuticulin). This structure was found to vary within the green elytra, where the reflector with 2 electron dense layers is most commonly seen comparing to the reflectors with 1 and 3 electron dense layers. We thus used the reflector with 2 electron dense layers to represent the green elytra. For modeling the layers, we repeatedly measured each layers including the top thick electron dense area at 26-30 points (green: 30 points, dark green: 27 points, and black: 26 points) along the elytral surface using ImageJ (Abràmoff et al., 2004). The lengths of the layers were subsequently used to calculate the expected reflection spectrum using reflectance calculator provided by Corporation Filmetrics®.

**Supplemental Figures**

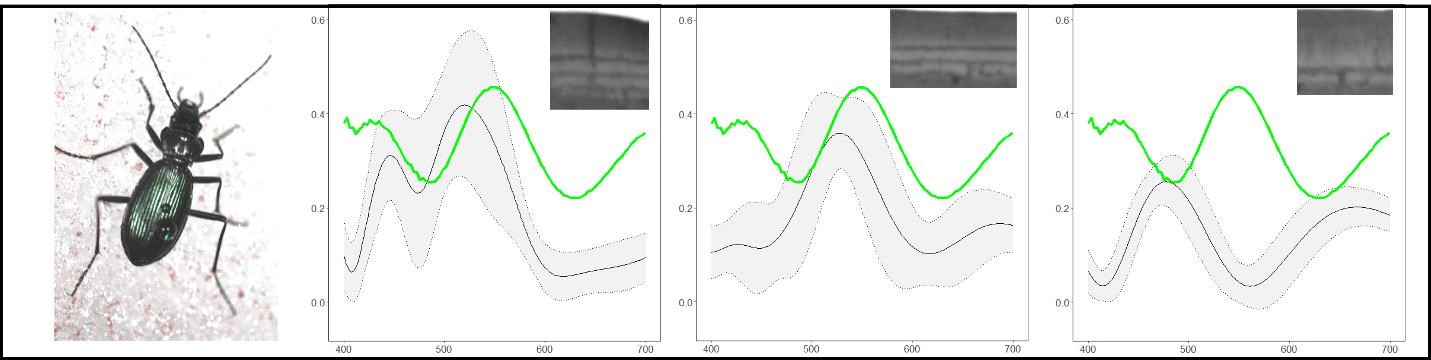

**Figure S1.** The calculated reflection spectra from different types of multilayer reflectors in the epicuticle. The beetle on the left shows the visual color of the elytra. Three types of multilayer reflectors (images on the top-right) with their calculated reflection spectrum (black lines) are shown with the observed spectrum (green line).
